# Robust calibration of hierarchical population models for heterogeneous cell populations

**DOI:** 10.1101/718270

**Authors:** Carolin Loos, Jan Hasenauer

## Abstract

Cellular heterogeneity is known to have important effects on signal processing and cellular decision making. To understand these processes, multiple classes of mathematical models have been introduced. The hierarchical population model builds a novel class which allows for the mechanistic description of heterogeneity and explicitly takes into account subpopulation structures. However, this model requires a parametric distribution assumption for the cell population and, so far, only the normal distribution has been employed. Here, we incorporate alternative distribution assumptions into the model, assess their robustness against outliers and evaluate their influence on the performance of model calibration in a simulation study and a real-world application example. We found that alternative distributions provide reliable parameter estimates even in the presence of outliers, and can in fact increase the convergence of model calibration.

**Highlights:**

- Generalizes hierarchical population model to various distribution assumptions
- Provides framework for efficient calibration of the hierarchical population model
- Simulation study and application to experimental data reveal improved robustness and optimization performance

## 1 Introduction

An important goal of systems biology is to obtain insights into the mechanisms and sources of cellular heterogeneity. This heterogeneity is critical for cellular decision making (Balázsi et al., 2011) and studied for various diseases and biological systems. To study heterogeneity, data at the single-cell level, e.g., time-lapse or snapshot data, are collected with experimental techniques such as fluorescent microscopy or flow cytometry.

To mechanistically study single-cell snapshot data, a variety of modeling approaches have been introduced. Ensemble models describe individual cells and model the overall population as a collection of many cells (Henson, 2003; Kuepfer et al., 2007). These models are often computationally expensive. To circumvent this, the cell population can be approximated by its statistical moments. Approximation approaches depend on assumptions about the contribution of intrinsic and extrinsic noise sources. While intrinsic noise is often defined as the stochasticity of gene expression, extrinsic noise is assumed to influence the reaction rates (Swain et al., 2002). Therefore, the statistical moments are obtained using the moment-closure approximation (Engblom, 2006), when intrinsic noise is assumed to be important, or Dirac-mixture approximations (Wang et al., 2019), when heterogeneity occurs mainly due to extrinsic noise. For the case of extrinsic noise, modeling approaches have been developed which infer the distribution of cellular properties using maximum entropy principles (Waldherr et al., 2009; Dixit et al., 2019). However, the aforementioned methods do not explicitly take into account subpopulation structures, which are omnipresent in heterogeneous cell populations (Altschuler and Wu, 2010). We recently introduced the hierarchical population model (Loos et al., 2018). This model describes subpopulation structures using mixture modeling, and ensures computational efficiency by employing approximations for statistical moments of the biological species of individual subpopulations. However, the model relies on a parametric assumption for the distribution of the subpopulations and only the multivariate normal distribution has been employed. This is a substantial limitation as the distribution of properties within subpopulations is often not normal (Pyne et al., 2009; Mar, 2019).

As the measured distribution reflects not only cellular heterogeneity, but also the variability of the measurement process, measurement noise and outliers might result in additional deviations from a normal distribution. In the analysis of ordinary differential equation (ODE) models, it has been shown that distributions with heavier tails than the normal distribution yield more robust parameter estimates in the presence of outliers (Maier et al., 2017). For single-cell snapshot data, the probability of observing outliers in the data is even higher due to the high number of data points (Pyne et al., 2009; Ilicic et al., 2016). Thus, heavy-tailed and skewed distributions are employed when studying flow cytometry (Pyne et al., 2009) or scRNA-seq data (Ding et al., 2019). However, the incorporation and assessment of distribution assumptions for the hierarchical population model is missing. Ideally, the employed distribution should not only provide reliable parameter estimates, but also yield a reasonable performance of model calibration.

In this manuscript, we describe the hierarchical population model and generalize it to different distribution assumptions. We derive the equations for the mathematical formulation of the model with alternative distribution assumptions. These equations, including the likelihood function and its gradient, are required to perform efficient model calibration using gradient-based maximum likelihood estimation. We analyze the influence of the distribution assumptions for simulated data of three biological motifs and three outlier scenarios. These outlier scenarios are motivated by experimental errors that can occur during data generation, e.g., due to a dropout event. Finally, we apply the hierarchical population model with the different distribution assumptions to analyze experimental data of NGF-induced Erk1/2 signaling.

## 2 Methods

### 2.1 Hierarchical population model

We consider single-cell snapshot data,

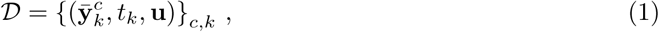

with indices *c* for the cell and *k* for the time point, stimulus vector 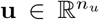 and a vector of measurements 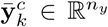 of the cell. The measured quantities might be, e.g., (relative) protein abundances. The measurement follows a distribution, which is a convolution of the measurement noise distribution and an additional distribution for a potential outlier-generating process. For a better readability, we neglected the index for varying stimuli.

In the hierarchical population model as introduced by Loos et al. (2018) the measured property of an individual cell is assumed to be distributed according to

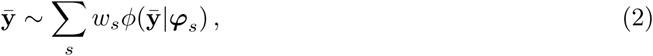

with relative subpopulation sizes *w*_*s*_ of subpopulations *s* = 1, …, *n*_*s*_ and parametric distribution *ϕ* which depends on the vector of distribution parameters ***φ***_*s*_ (Fig. 1A). The density *ϕ* captures biological variability due to intrinsic or extrinsic noise as well as measurement noise and the outlier distribution. The relative subpopulation sizes *w*_*s*_ are positive and sum up to one. The distribution parameters ***φ***_*s*_ arise from the single-cell dynamics and models for cell-to-cell variability. Individual single-cell trajectories can be obtained using Markov jump processes (Gillespie, 1977), ordinary (Klipp et al., 2005) or stochastic differential equations (Gillespie, 2000). Instead of simulating the cells individually, which can become time-consuming, we compute the temporal evolution of the statistical moments, i.e., means 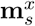 and covariances 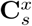, of biochemical species **x**. The temporal evolution is defined by a function *g*_*z*_, e.g., obtained by moment-closure (Engblom, 2006), sigma-point (van der Merwe, 2004) or Dirac mixture approximations (Wang et al., 2019), depending on the assumptions about intrinsic and extrinsic noise. This yields

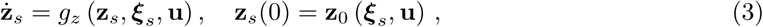

with 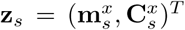, initial conditions **z**_0_ and subpopulation parameters ***ξ***_*s*_ = (***β***_*s*_, ***D***_*s*_), with 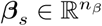 and 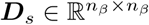. These parameters are given by

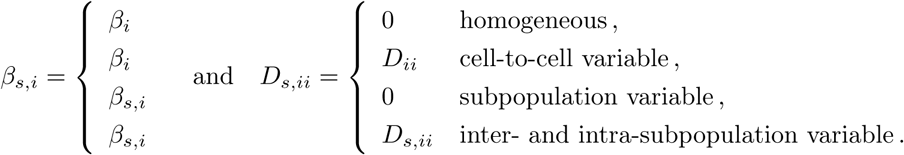

in which *β*_*s,i*_ denotes the *i*th element of ***β***_*s*_, and *D*_*s,ii*_ denotes the *i*th diagonal element of ***D***_*s*_. The parameter *β*_*s,i*_ encodes the mean of the cellular property, while the parameter *D*_*s,ii*_ encodes its spread within a subpopulation. Homogeneous parameters are assumed to be the same for all cells of the whole cell population; cell-to-cell variable parameters differ between cells of the same subpopulation; subpopulation variable parameters differ between but not within a subpopulation; and inter- and intra-subpopulation variable parameters differ both between and within subpopulations (Fig. 1B).

**Figure 1:**
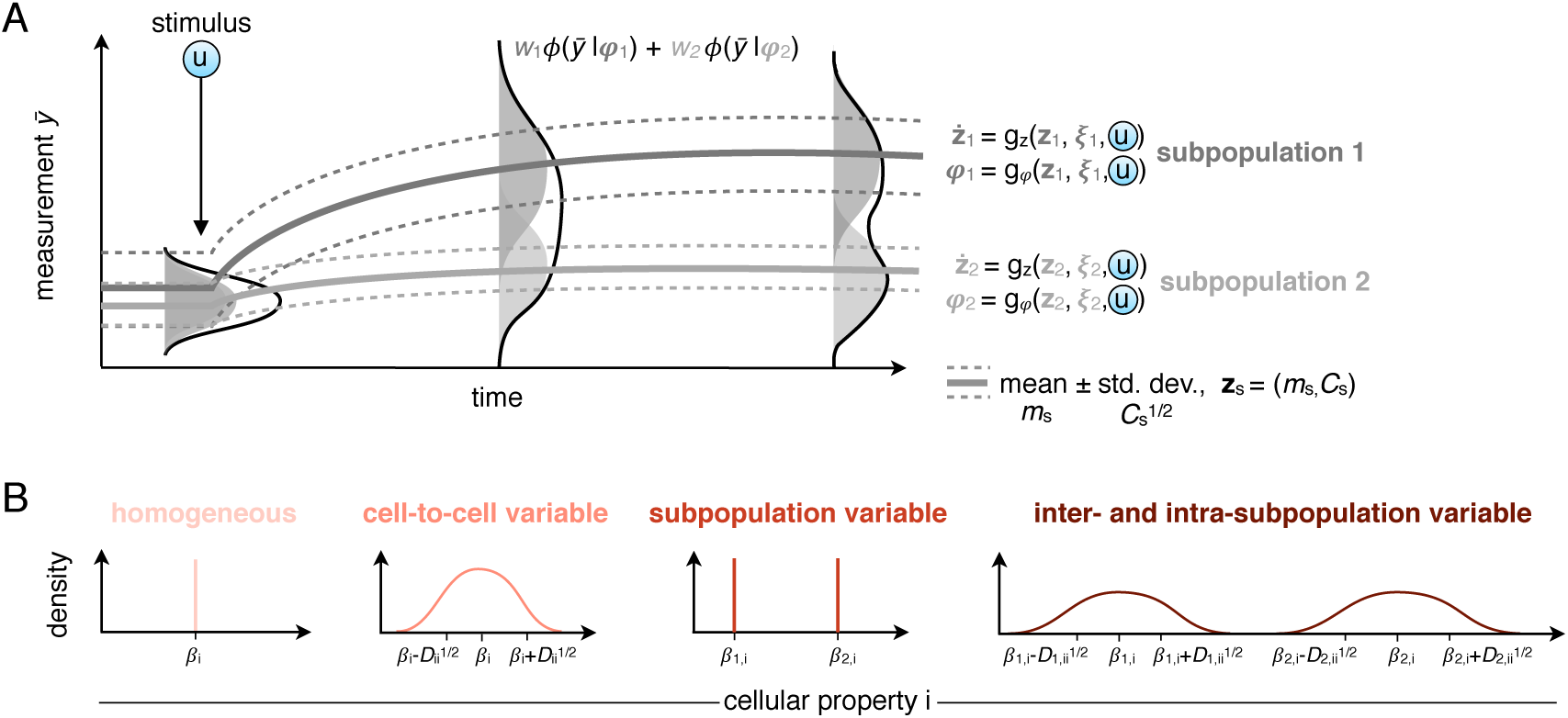
Illustration of the hierarchical population model. (A) The cell population comprises two heterogenenous subpopulation which respond differently to stimulation. Means and covariances of the species for each subpopulation are linked to a mixture distribution. The light and dark gray lines show the means of the subpopulations and the black line shows the distribution of the whole cell population. (B) Heterogeneity is captured by assuming each parameter/cellular property to be distributed according to one of the indicated cases.

The moments for a subpopulation are mapped to the distribution parameters,

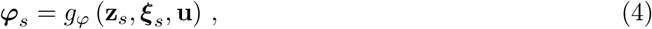

of a distribution *ϕ*. Thus, *g*_*φ*_ encodes the mapping from the biochemical species to the observables, i.e., the measurable output of the system, as well as the mapping of the observables to the distribution parameters (see Appendix D.1 for an example). So far, the multivariate normal distribution has been employed,

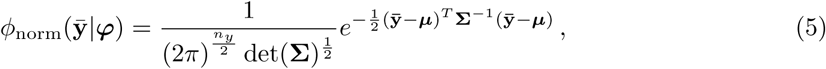

with distribution parameters ***φ*** = (***μ*, Σ**), comprising mean 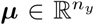 and covariance matrix 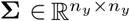. This yields

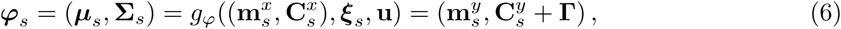

including measurement noise **Γ**, which is generally assumed to be the same for all subpopulations.

#### 2.1.1 Calibration of the hierarchical population model

The parameters of the model, including relative subpopulation sizes *w*_*s*_, subpopulation parameters ***ξ***_*s*_ and parameters for the measurement noise need to be estimated from data. For this, we denote the overall parameter object by ***θ*** (see Appendix D.1 for an example). The model is calibrated by maximizing the likelihood function,

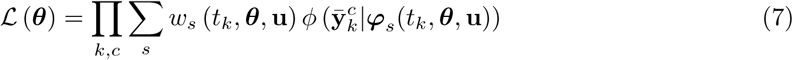

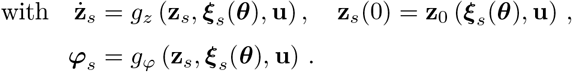

This can efficiently be done by multi-start local optimization employing the gradient of the likelihood function (Raue et al., 2013; Loos et al., 2016). In the following, we provide the likelihood functions for several distribution assumptions which can be incorporated into the hierarchical population model. For each distribution, we derive the function *g*_*φ*_ introduced in (4), which maps the mean and covariances of the species to the distribution parameters ***φ***_*s*_.

### 2.2 Alternative distribution assumptions for the hierarchical population model

We considered two alternatives to the normal distribution: the skew normal and the Student’s t distribution (Fig. 2) (see Appendix B for a third distribution, the negative binomial distribution). In the following, we discuss these distributions and provide the equations which are required to incorporate the distributions in the hierarchical population model.

**Figure 2:**
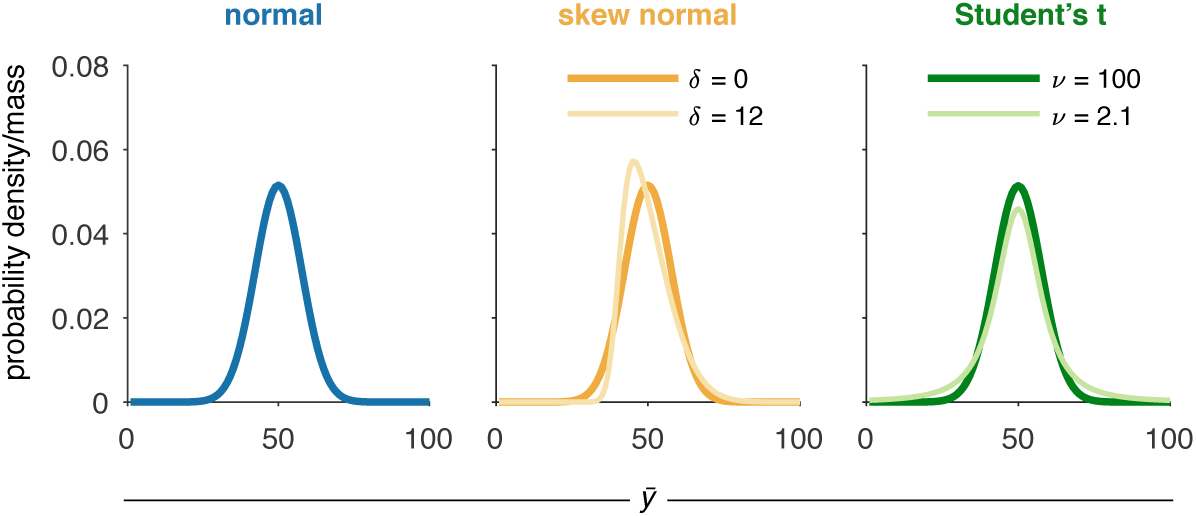
Distributions assumptions for the hierarchical population model. The visualized distributions (normal, skew normal and Student’s t) for 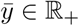 have mean *m* = 50 and variance *C* = 60. The skew normal and Student’s t distributions are visualized for different skewness parameters *δ* and degree of freedom *ν*, respectively.

#### 2.2.1 Multivariate skew normal distribution

A challenge in the analysis of single-cell data is that the observed cell population is often skewed (Pyne et al., 2009). Therefore, distributions which account for skewness are often employed in the analysis of, e.g., flow cytometry data (Johnsson et al., 2016). The multivariate skew normal distribution has distribution parameters ***φ*** = (***μ*, Σ, *δ***), with location 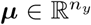, covariance matrix 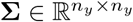 and skew parameter 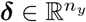. The probability density function is

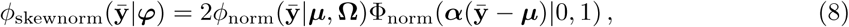

with 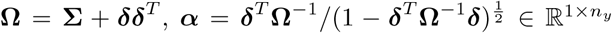 and Φ_norm_ denoting the cumulative distribution function of a univariate standard normal distribution. If ***δ*** = **0**, the distribution equals a multivariate normal distribution. As provided by Pyne et al. (2009) and Sahu et al. (2003), the mean and covariance matrix of the multivariate skew normal distribution are

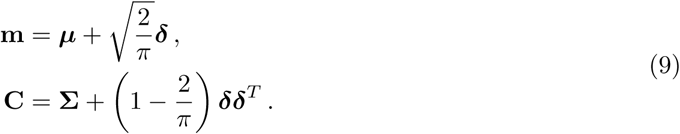

This yields for ***φ***_*s*_(***θ***) = (***μ***_*s*_(***θ***), **Σ**_*s*_(***θ***), ***δ***(***θ***)) the relation

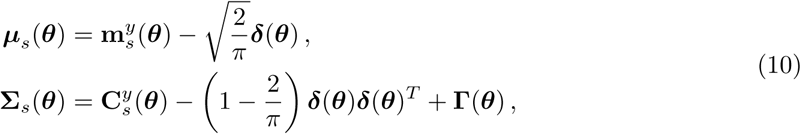

with measurement noise matrix **Γ**. The entries of the skew parameter vector ***δ*** are allowed to differ in each dimension. However, they are restricted in a way that **Σ**_*s*_ needs to be positive definite. The derivatives of the probability density (8) and the distribution parameters (10) are provided in Appendix A.2.

#### 2.2.2 Multivariate Student’s t distribution

The Student’s t distribution is often employed as a robust alternative to the normal distribution in regression (Lange et al., 1989), modeling of population-average data (Maier et al., 2017), and in the analysis of single-cell data (Lo et al., 2008; Pyne et al., 2009; Ding et al., 2019). The tails of the Student’s t distribution are heavier than the tails of a normal distribution, and thus the distribution can better cope with outliers in the data. The multivariate Student’s t distribution has distribution parameters ***φ*** = (***μ*, Σ**, *ν*) with location 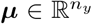, shape matrix 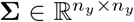 and degree of freedom *ν* ∈ ℝ_+_. The probability density function reads

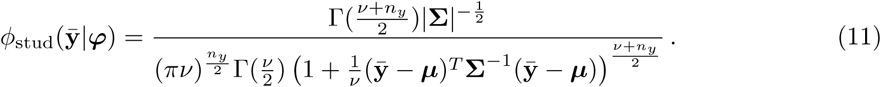

For *ν* > 2, the mean and covariance matrix of the multivariate Student’s t distribution are given by

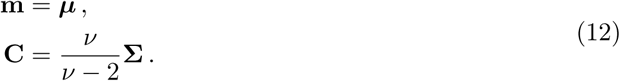

This yields ***φ***_*s*_(***θ***) = (***μ***_*s*_(***θ***), **Σ**_*s*_(***θ***), *ν*(***θ***)) with

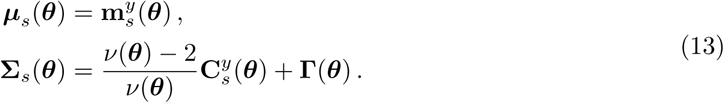

For *ν* → ∞, the Student’s t distribution equals a normal distribution. The derivatives of the probability density (11) and the distribution parameters (13) are provided in Appendix A.3.

## 3 Results

### 3.1 Evaluation of influence of distribution assumptions for simulated data

To assess the performance of model calibration under the different distribution assumptions in the hierarchical population model, we simulated data for three different motifs: a conversion process, a two-stage gene expression and a birth-death process (Fig. 3 and Appendix D). The first motif is frequently found in signal transduction networks, and the last two motifs models are commonly used for the description of gene expression. In this study, we assumed that the cell population comprises two subpopulations and that the true underlying difference between the subpopulations is known in the hierarchical population models. However, the hierarchical population model is able to describe more than two subpopulations and the number of subpopulations could be inferred by performing model selection.

**Figure 3:**
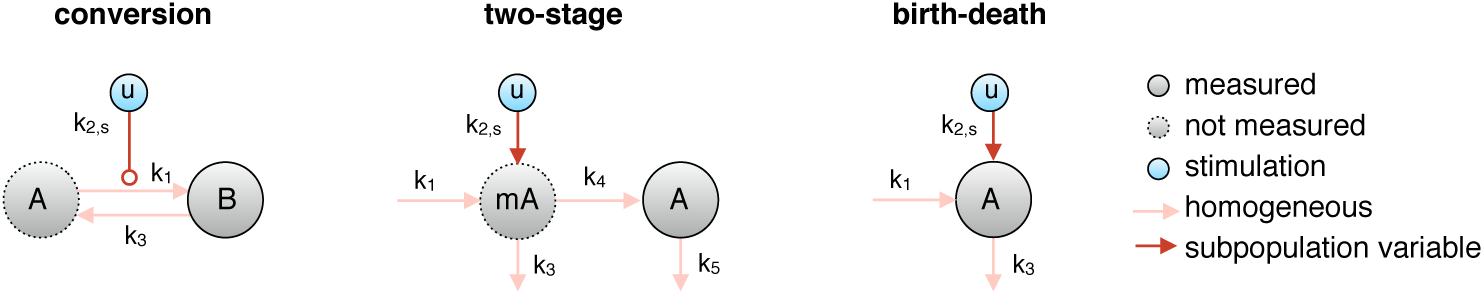
Three motifs for the simulation study. We considered a conversion process, a two-stage gene expression and a birth-death process. For each motif, we assumed two subpopulations which differ in their response to stimulus *u*. All other reaction rate constants were assumed to be homogeneous, i.e., the same for all cells.

All systems were assumed to be in steady state before the stimulus **u** was added at time point 0. However, this is not required by the hierarchical population model. For each motif, we chose three parameter vectors, three numbers of time points and four numbers of cells per time point (50, 100, 500, 1000). This yielded 108 data sets which were simulated using the stochastic simulation algorithm (Gillespie, 1977). The differences in the measurements for cells within a subpopulation arose solely due to intrinsic noise and no additional measurement noise was added to the data.

We perturbed the data according to different outlier scenarios (Fig. 4):

- *No outliers*: no outliers were included in the data.
- *Zeros*: the measured concentration at a certain time point *t*_*k*_ is zero, e.g., due to a missing label or entry or a dropout event (Luecken and Theis, 2019). Consequently, we measured 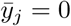 for randomly chosen cells.
- *Doublets*: one measurement includes the summed information of two cells, e.g., due to wrongly measuring two cells instead of one. As a simplified simulation approach, the measured value of randomly chosen cell was doubled.
- *Uniform*: the measurement does not carry any information. Randomly chosen cells have uniformly distributed values in a defined regime (C.21) instead of the real measurement.

**Figure 4:**
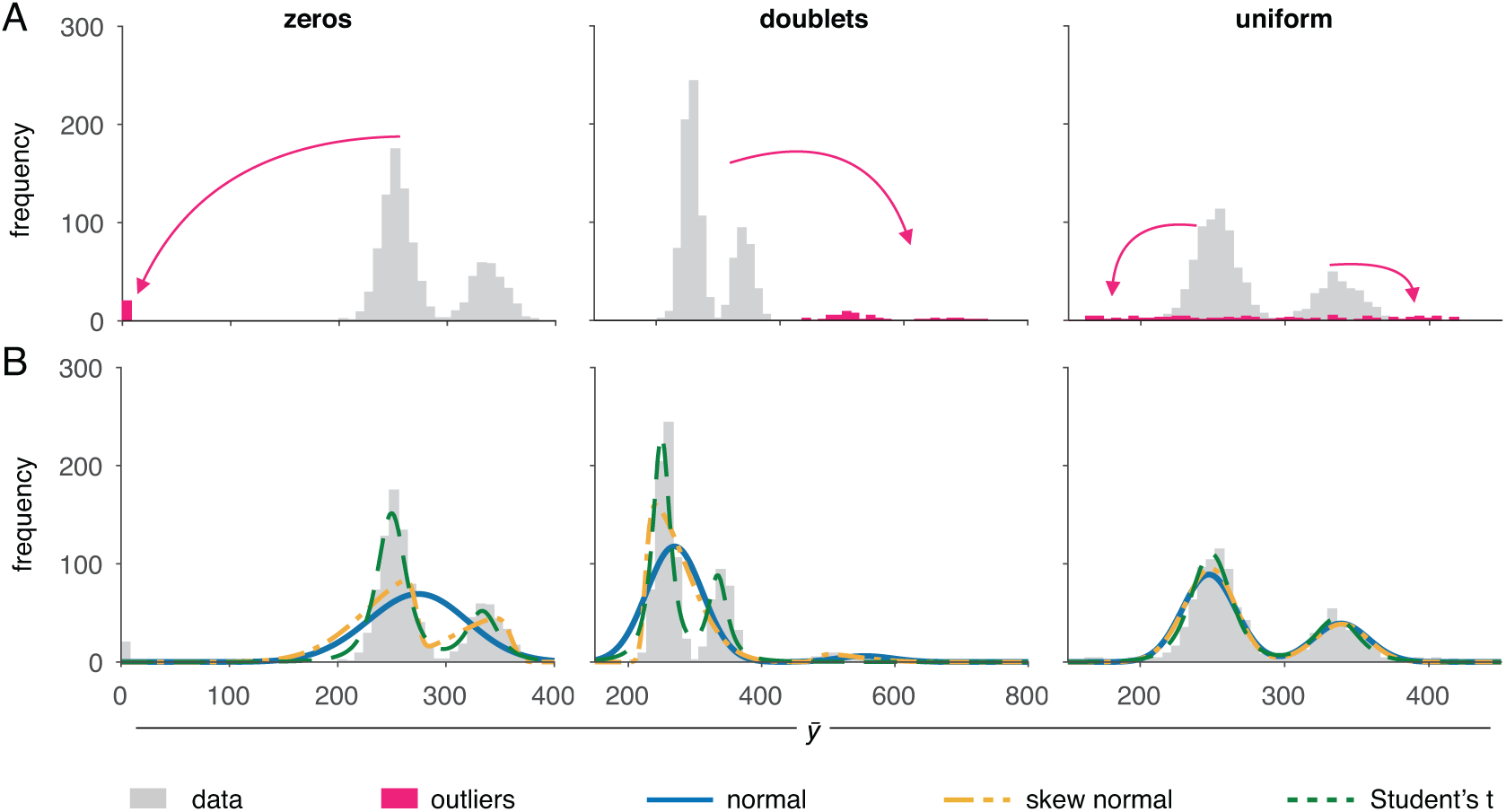
Outlier scenarios for single-cell snapshot data. (A) Example data sets of a conversion process with (B) corresponding fits with different distribution assumptions. The arrows in (A) illustrate the outlier-generating mechanisms.

In this simulation study, the deviation between the biological quantity and the mean of the quantity for a subpopulation arises due to intrinsic noise. The discrepancy between the true biological quantity and the measurement only arises due to the outlier-generating process. The amount of outliers in the data has a different influence for the different scenarios. The introduction of *zero* measurements is, for example, a bigger perturbation of the data as the *uniform* scenario. To obtain a comparable perturbation, we used different percentages of outliers for the scenarios and assumed 2%, 5% and 10% of the cells to be outliers for scenarios *zeros, doublets*, and *uniform*, respectively.

We calibrated the hierarchical population models based on all data sets for the different distribution assumptions with 30 optimization runs which were started at randomly chosen parameter values. For this simulation study, the measurements were count data and the continuous distributions were evaluated for the untransformed measured counts. The subpopulation sizes were fitted in linear space and the parameters required for the simulation of the statistical moments in log_10_ space. We assumed the true underlying sources of heterogeneity to be known and allowed for measurement noise. The moments of the subpopulations were obtained using the moment-closure approximation.

The fits for the different distribution assumptions are shown for three example data sets in Fig. 4B.

All distributions accurately fitted the data for the *uniform* outlier scenario. However, for the *zeros* and *doublets* scenarios, only the Student’s t distribution was not deviated by the outliers, potentially because it has the heaviest tails of the considered distributions.

In the following comparison of the distributions, we only distinguished between the motifs and the absence or presence of outliers. The data sets corresponding to different parameter values, time points, number of cells and outlier scenarios were merged. The comparison of the individual outlier scenarios is displayed in Appendix Fig. D1.

To compare the different models quantitatively, we used the Bayesian Information Criterion (BIC) (Schwarz, 1978),

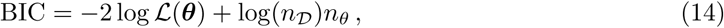

with *n*_𝒟_ denoting the number of data points and *n*_*θ*_ denoting the number of parameters. As lower BIC values are preferable (Schwarz, 1978), the BIC rewards high likelihood values and penalizes model complexity. As an alternative, the Akaike Information Criterion (AIC) could be used (Akaike, 1973). We compared the differences in BIC values to the minimal BIC value found for the given data set (=ΔBIC) (Fig. 5A). In the case of outlier-free data, the best distribution assumption differed between the studied motifs. For the conversion process the normal distribution was most appropriate, while for the two-stage gene expression and the birth-death motif the normal and the skew normal distribution achieved low BIC values. As soon as outliers were introduced to the data, the Student’s t distribution provided in average the lowest BIC value for all considered motifs and was therefore selected as most suited model.

**Figure 5:**
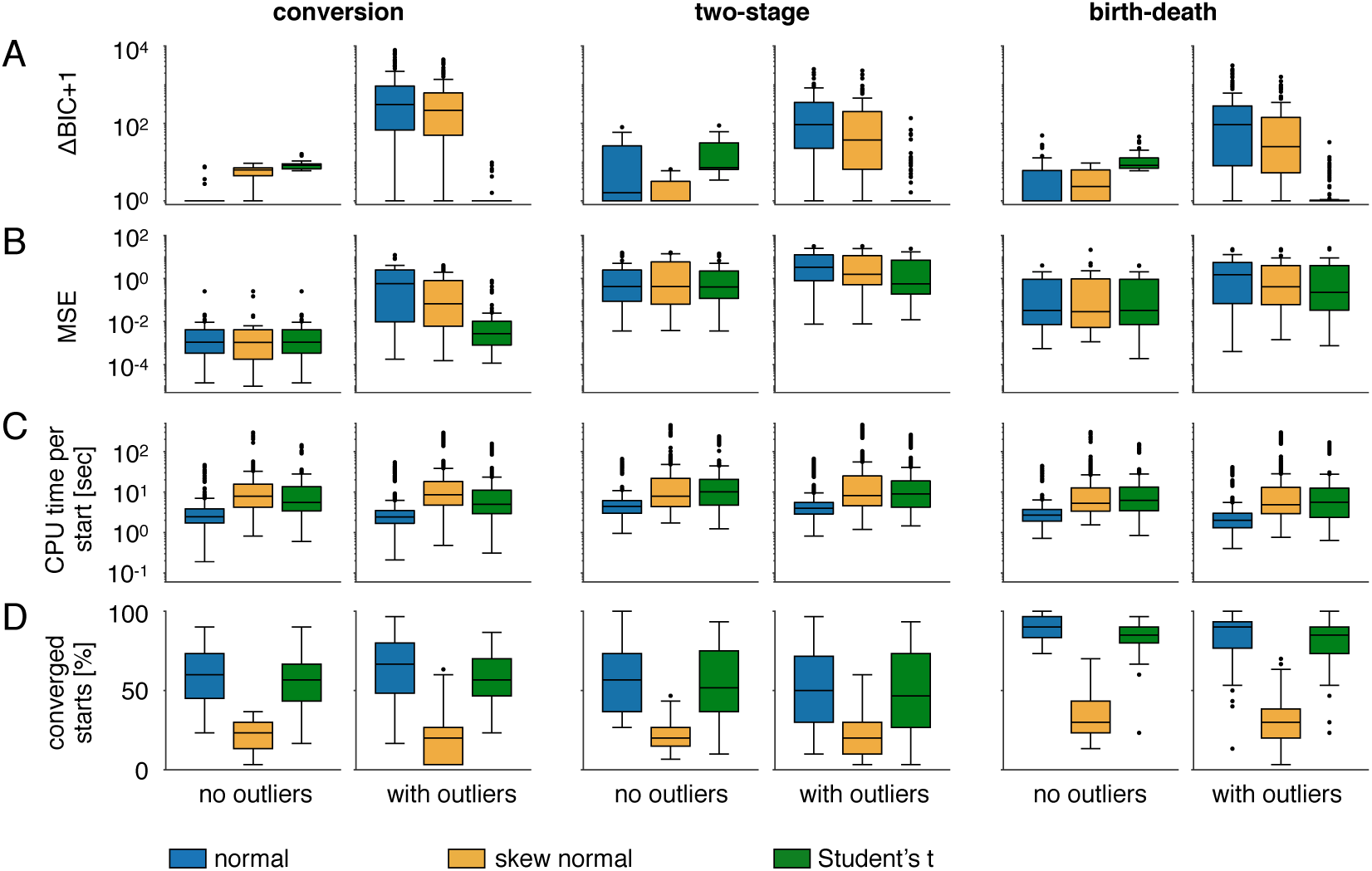
Results for the simulation study. Comparison of (A) ΔBIC values, (B) MSE, (C) CPU time per optimization start and (D) number of converged starts for the distribution assumptions for the motifs conversion process, two-stage gene expression and birth-death process. We considered outlier-free and outlier-corrupted data. For *no outliers*, each boxplot in (C) has 1080 points (36 data sets and 30 optimization runs) and the other boxplots (A, B, D) comprise 36 points. The subplots for *with outliers* comprise the results of all three outlier scenarios. For *with outliers*, each boxplot in (C) has 3240 points (3 outlier scenarios, 36 data sets, 30 optimization runs) and the other boxplots (A, B, D) comprise 108 points.

The mean squared error (MSE) provides a measure for the accuracy of the parameter estimates:

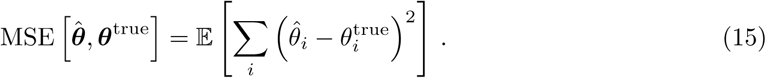

A small (norm of the) MSE indicates a good agreement of the true and estimated parameters. A high MSE indicates a large difference to the true parameters, which in turn might distort model predictions and the capability of the model to provide correct insights into the underlying system of study. We considered the MSE in the logarithmic/linear space in which the parameters were fitted and summed the errors of the parameter estimates to obtain a single value for each data set and distribution. The true parameters which were used to generate the data for the simulation study are known, and we compared the MSEs of the parameter estimates obtained with the different models (Fig. 5B). For the cases of *no-outliers*, the MSE for all distributions were similar. In the presence of outliers, the Student’s t distribution still provided low MSEs which were comparable to the those obtained in the outlier-free scenario. The MSE obtained by the normal and skew normal distribution increased in the presence of outliers. The skew normal distribution provided slightly more robust results, i.e., lower MSEs, than the normal distribution. High MSEs do not necessarily result in deviations of the output of the model, e.g., if the output is insensitive to these parameters. Thus, we compared the model outputs for the parameters which were obtained for the outlier-corrupted data to the original, *no-outlier* data (Appendix Fig. D1E). We found that the Student’s t distribution consistently outperformed the other distributions.

A further important aspect to consider is the robustness and efficiency of model calibration. Often many models which represent different biological hypotheses need to be calibrated. These hypotheses could include different sources of heterogeneity or different numbers of subpopulations. Therefore, the optimization of each individual model should be fast and robust. To assess the performance of model calibration using the distribution assumptions, we considered the computation time and number of converged starts (Fig. 5C-D). We considered an optimizer start to be converged, if the difference between the obtained log-likelihood value and the best log-likelihood function for this distribution assumption and the considered motif is below 10^−3^. For most of the here considered motifs and data sets the computation time and convergence were not substantially influenced by the presence of outliers. On average, the normal distribution required the lowest computation time. An explanation for this could be that the optimization problem using this distribution is lower than the problem using the skew normal and the Student’s t distribution, since no additional parameters were estimated from the data. In terms of converged starts, we observed some differences between the considered motifs. For all motifs and scenarios the skew normal distribution provided the lowest number of converged starts, while the normal and Student’s t distribution did not suffer from convergence problems.

The simulation study showed that the consideration of alternative distribution assumptions was beneficial and the normal distribution was often outperformed by the other distributions. The heavier tails of the distributions allowed for a compensation of the outliers in the data, but did not generate substantial computational overhead in the outlier-free case. Overall, the Student’s t distribution provided reliable parameter estimates, the overall closest predictions to the original *no-outlier* data and, at the same time, enabled an efficient calibration of the corresponding model.

### 3.2 NGF-induced Erk1/2 signaling

To test the distributions in a real application setting, we reanalyzed the data and hierarchical population model studied in (Loos et al., 2018). The model describes binding of NGF to its receptor TrkA, which then induces the phosphorylation of Erk1/2 (Fig. 6A), an important process involved in pain sensitization. The measurements were obtained from primary sensory neurons using fluorescent microscopy. For more details on the model and data, we refer to Loos et al. (2018).

**Figure 6:**
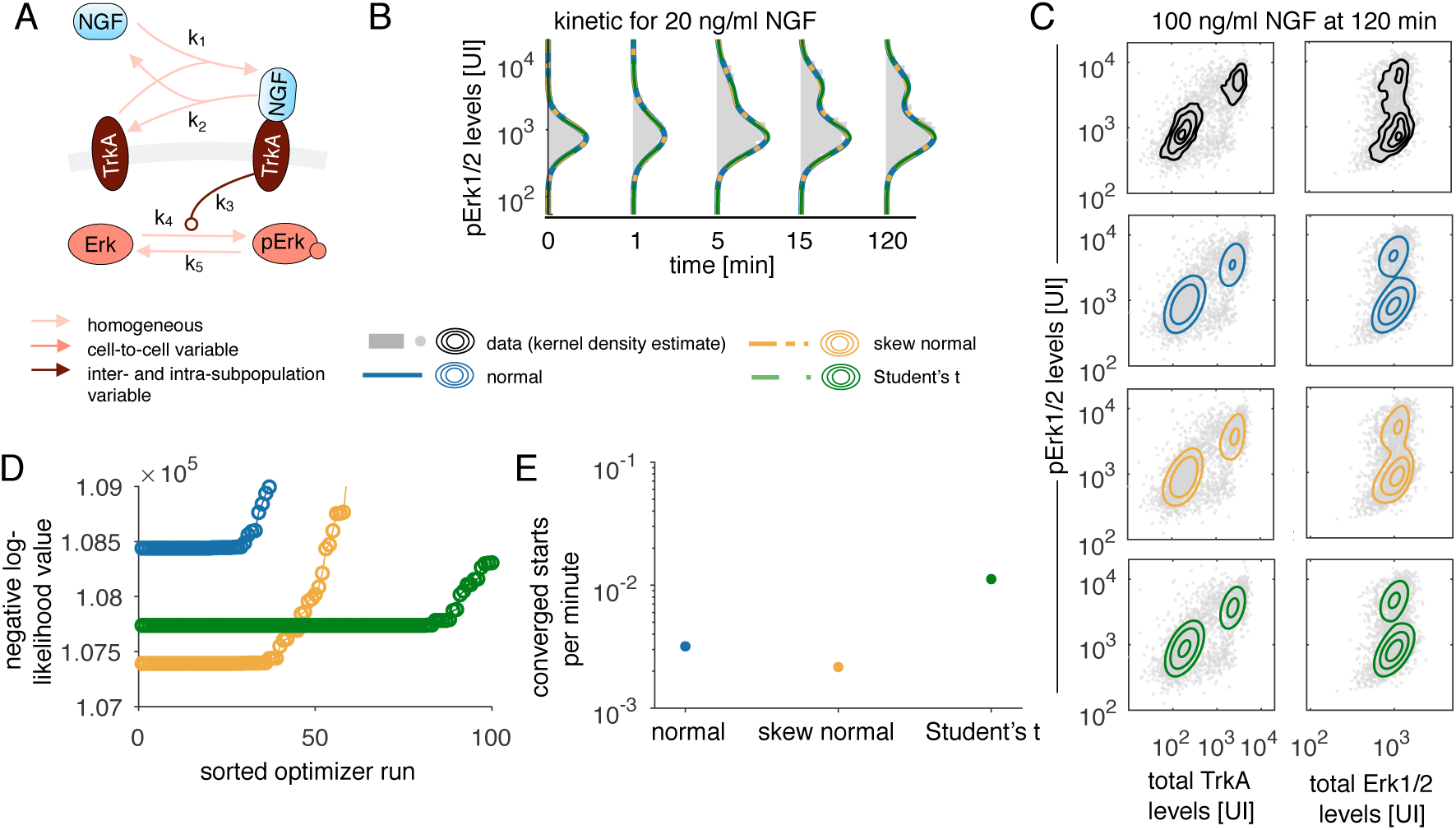
Robust distributions for NGF-induced Erk1/2 signaling. (A) Model for NGF-induced Erk1/2 signaling. (B,C) Data and model fits for (A) univariate measurements of pErk1/2 levels and (C) bivariate measurements for pErk/TrkA and pErk/Erk levels. (D) Likelihood waterfall plot for the three different distribution assumptions. The best 80 values are shown and in total 500 optimization runs started at randomly drawn parameter values were performed. (E) The performance of the optimization measured as number of converged starts per minute.

For this analysis, we log-transformed the simulation and data. For the Student’s t distribution we assumed that one parameter for the degree of freedom was shared across the dimensions, yielding one additional parameter. For the skew normal distribution, each dimension is allowed to have different skewness parameters, yielding two parameters more than the normal distribution. Calibrating the hierarchical population models, we found that for the univariate data of the pErk1/2 kinetics, the model fits cannot visually be distinguished (Fig. 6B and Fig. D2). The skew normal distribution fitted the bivariate data the best (Fig. 6C,D and Fig. D2). We could not assess the MSE since the true parameters are not known. Visualizing the likelihood waterfall plots (Fig. 6D) and analyzing the performance of the optimization (Fig. 6E), we found that the Student’s t distribution substantially outperformed the other distributions in terms of converged optimizer starts (per minute). Interestingly, the skew normal which showed bad convergence for the simulation study even had a higher convergence than the normal distribution for this application problem. The skew normal provided the best likelihood and BIC value. An explanation for why the skew normal distribution was chosen over the other distributions could be a skewed distribution of Erk levels arising from biological variability rather than noise or outliers. However, the hierarchical population model uses a single distribution assumption for the combined influence of cell-to-cell variability, measurement noise and outliers. Accordingly, if the distribution assumptions allow for skewness, the skewness can arise from any of the properties. To summarize, the incorporation of the alternative distribution assumptions into the hierarchical population model demonstrated their robustness and efficiency not only for the simulation study, but also for the considered experimental data set. Thus, the distribution assumptions seem also promising for real experimental data and future work should include the evaluation of the distributions for more experimental data.

## 4 Discussion

The hierarchical population model is a suitable tool for studying cellular heterogeneity, but it requires appropriate distribution assumptions for the cellular subpopulations. Here, we incorporated various distributions for the subpopulations and provided the equations to perform gradient-based model calibration. Gradient-based calibration has previously has been shown to be highly efficient for the considered model class (e.g., Loos et al. (2016)) and, thus, also enhances optimization-based uncertainty analysis using profile likelihoods (Raue et al., 2009). This efficiency of model calibration is especially important when a large number of hierarchical population models, which represent different hypothesis such as differences between or number of subpopulations, needs to be compared. Furthermore, we studied the influence of the choice of distribution on the estimation results and the performance of model calibration for artificial and real experimental data.

Differences in measurements of single-cell arise due to various factors: biological variability, measurement noise and the outlier-generating mechanisms. The distribution incorporated in the hierarchical population model ideally should capture all these factors. Here, we found that the normal distribution assumption is often appropriate when the biological variability yields a normal distribution of the subpopulations and additionally a limited number of outliers is to be expected. However, the biological variability might not always yield a normal distribution of the subpopulations. This was observed in the experimental data of the primary sensory neurons and suggests that the best choice of distribution highly depends on the particular application problem. This motivates the use and comparison of alternative distributions also when no outliers are present in the data. If the data is outlier-corrupted, alternatives such as the skew normal or Student’s t distribution are more reasonable. The Student’s t distribution provided reliable results when the data is outlier-free and could be considered as default distribution assumption. If more information is available about the precise type of outliers, e.g., if they arise due to dropout events in scRNA-seq data, computational methods can be adapted accordingly (Pierson and Yau, 2015; Eraslan et al., 2019). While the Student’s t distribution suffered from problems of over-fitting in the case of population-average data (Maier et al., 2017), the number of measurements in single-cell data sets is usually much higher and we do not expect to face the same problems as for population-average data.

Other distributions could be incorporated into the hierarchical population model, given that their mean and covariance are finite and an analytical gradient can be provided. To allow for different degrees of freedom in multivariate measurements, a t copula could be employed (Luo and Shevchenko, 2010). Also, a skewed version of the Student’s t distribution as, e.g., used by Pyne et al. (2009) could be incorporated. The log-normal distribution could easily be incorporated by transforming the observable. In this case, a constant factor needs to be added to the BIC value when performing model selection.

In summary, we introduced the use of different distribution assumptions for the hierarchical population model and evaluated their performance. Our results on simulation and application examples suggested that these distribution assumptions substantially improve the hierarchical population model and the reliability of its results, and, thus, enhance the study of cellular heterogeneity.

## Implementation and code availability

The alternative distribution assumptions are implemented in the MATLAB toolbox ODEMM (Loos et al., 2018) available under https://github.com/ICB-DCM/ODEMM. The model calibration was performed using the toolbox PESTO (Stapor et al., 2018). The ODE models were simulated using the toolbox AMICI (Fröhlich et al., 2017). The sigma-point approximation was obtained by the SPToolbox. These toolboxes can be found under https://github.com/ICB-DCM. For the momentclosure approximation, we employed the toolbox CERENA (Kazeroonian et al., 2016) available under https://cerenadevelopers.github.io/CERENA/. The whole analysis was performed using MATLAB 2017b. The code to reproduce the results of this study is available under http://doi.org/10.5281/zenodo.3354136.

## Acknowledgements

This work was supported by the German Ministry of Education and Research by the grant SYS-Stomach (01ZX1310B) and the grant INCOME (FKZ 01ZX1705A).

## A Gradient of the likelihood functions

### A.1 Multivariate normal distribution

The probability density function for the multivariate normal distribution is defined in (5). The log-density function is given by

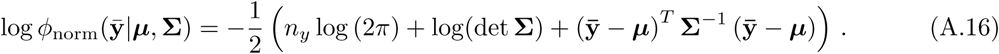

Assuming that the distribution parameters depend on parameter vector ***θ***, the derivative of the multivariate normal density is given by

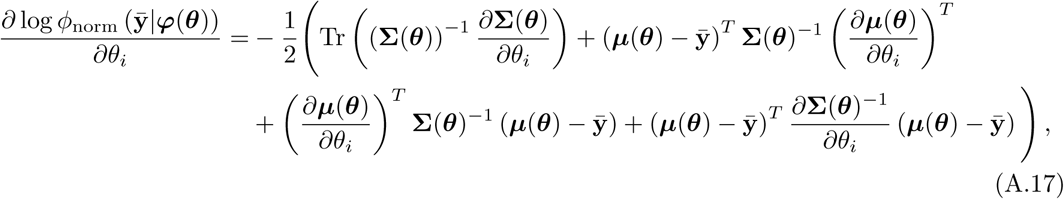

with derivatives for the distribution parameters for subpopulation *s*

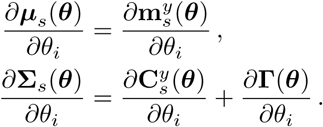

### A.2 Multivariate skew normal distribution

The probability density function for the multivariate skew normal distribution is defined in (8). The log-density function is given by

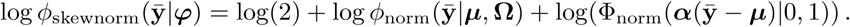

Assuming that the distribution parameters depend on parameter vector ***θ***, the gradient is given by

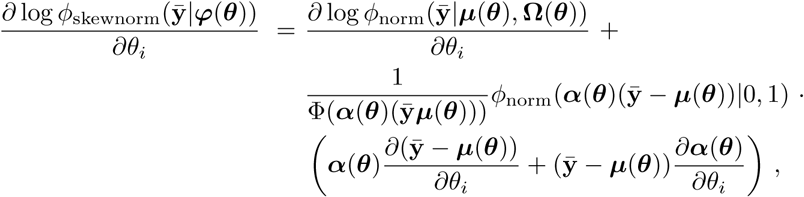

with

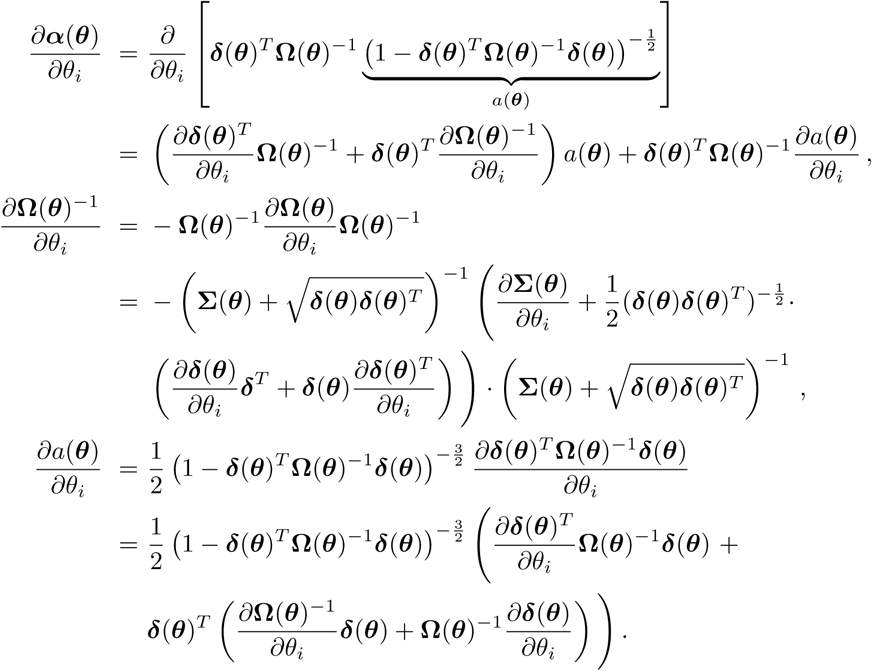

The derivatives for the distribution parameters (10) for subpopulation *s* are given by

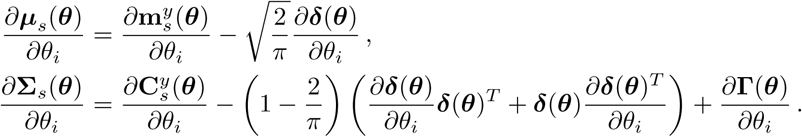

### A.3 Multivariate Student’s t distribution

The probability density function for the multivariate Student’s t distribution is defined in (11).

The log-density function is

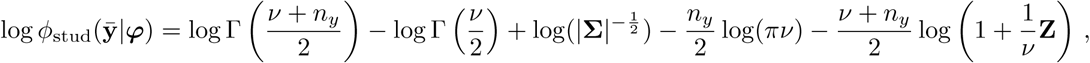

with 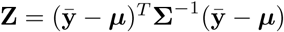. The gradient is given by

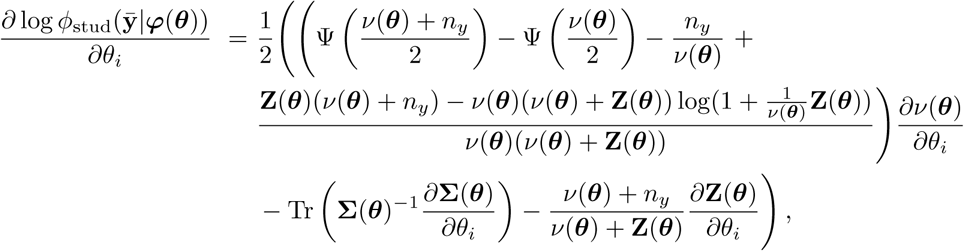

with

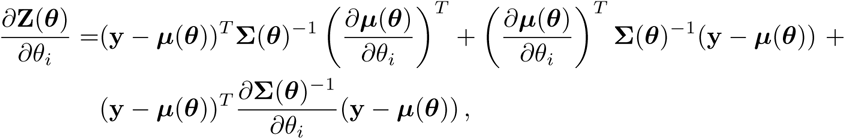

and digamma function denoted by Ψ.

The derivatives for the distribution parameters (13) for subpopulation *s* are

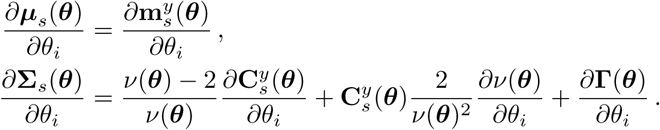

## B Negative binomial distribution

A further distribution assumption which is often employed in the analysis of single-cell data is the negative binomial distribution (Grün et al., 2014; Amrhein et al., 2019), which is a count distribution in contrast to the other distributions. For a two-stage model of gene expression, the protein number in steady state follows a negative binomial distribution if the ratio of mRNA degradation to protein degradation is high (Shahrezaei and Swain, 2008), or if the mRNA molecules are produced in bursts (Amrhein et al., 2019). This distribution has the parameters ***φ*** = (*τ, ρ*) with *τ* > 0 and *ρ* ∈ [0, 1]. We considered the univariate case and multivariate data could be modeled by the product distribution which neglects correlations. The probability mass function reads

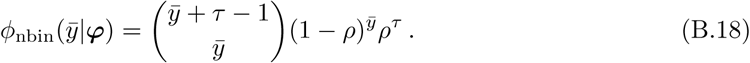

The mean and variance of the negative binomial distribution are

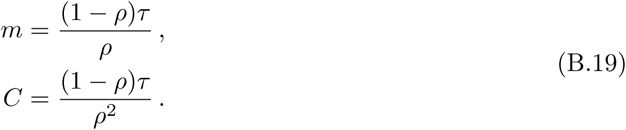

Thus, the distribution parameters ***φ***_*s*_(***θ***) = (*ρ*_*s*_(***θ***), *τ*_*s*_(***θ***)) are mapped to the moments via

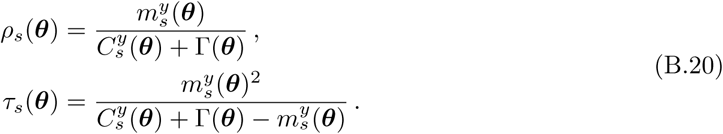

The derivatives of the probability density (B.18) and the distribution parameters (B.20) are provided in Appendix B. For large *τ*, the negative binomial distribution approaches a normal distribution.

The log-density function reads

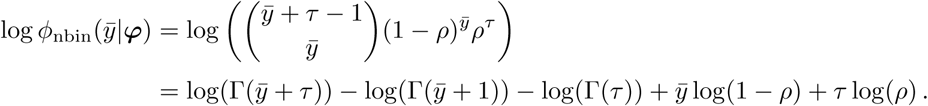

The derivative of the log-density function is

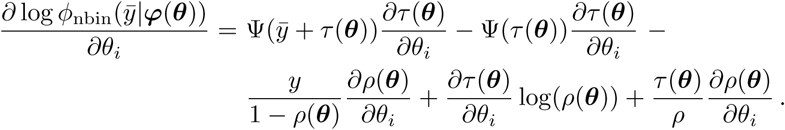

The derivatives of the distribution parameters (B.20) for subpopulation *s* are

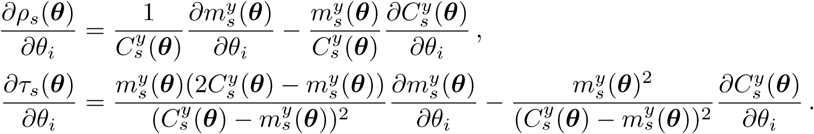

In contrast to the other distributions, the negative binomial distribution can not take into account correlation structures between the dimensions of multivariate data. A multivariate extensions which account for correlations was proposed by Shi and Valdez (2014).

## C Uniform outlier scenario

In the *uniform* outlier scenario, the measured values of the outlier cells were assigned to the rounded value of uniformly distributed values on the interval

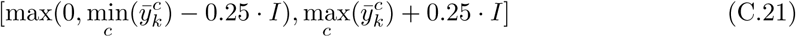

with *I* denoting the length of the interval without outliers.

## D Models

### D.1 Conversion process

The conversion process is described by the following reactions:

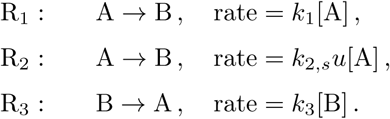

Reaction R_1_ describes the basal conversion from A to B and reaction R_2_ the stimulus-dependent conversion. The conversion from B to A, reaction R_3_, does not depend on stimulus *u*. We assumed mass conservation with [A] + [B] = 1000.

The moment-closure approximation provides the temporal evolution of the moments,

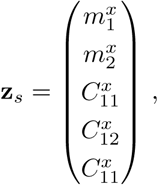

with 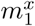 denoting the mean of species A, 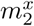 denoting the mean of species B and 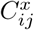 denoting the corresponding entries of the covariance matrix. The measurement noise is defined as **Γ** = *Σ*_noise_. The subpopulation parameter vector for subpopulation *s* = 1, 2 is given by ***ξ***_*s*_ = (***β***_*s*_, **D**_*s*_) with

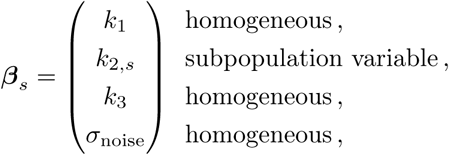

and **D**_*s*_ = **0**.

The mapping *g*_*φ*_ introduced in (4) then encodes the mapping from the biochemical species A and B to its observable B, as well as the mapping to the distribution parameters as in (6, 10, 13, B.20). The overall parameter object which is estimated from the data is given by

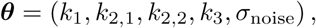

for the normal distribution,

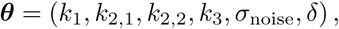

for the skew normal distribution, and

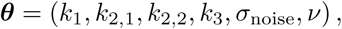

for the Student’s t distribution,

### D.2 Two-stage gene expression

The two-stage gene expression is described by the following reactions:

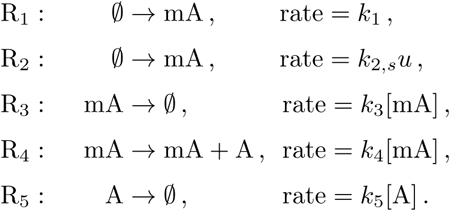

Reactions R_1_ and R_2_ describe the stimulus-independent and stimulus-dependent mRNA expression, respectively. Reaction R_3_ describes mRNA degradation, reaction R_4_ protein expression and reaction R_5_ protein degradation.

**Figure D1:**
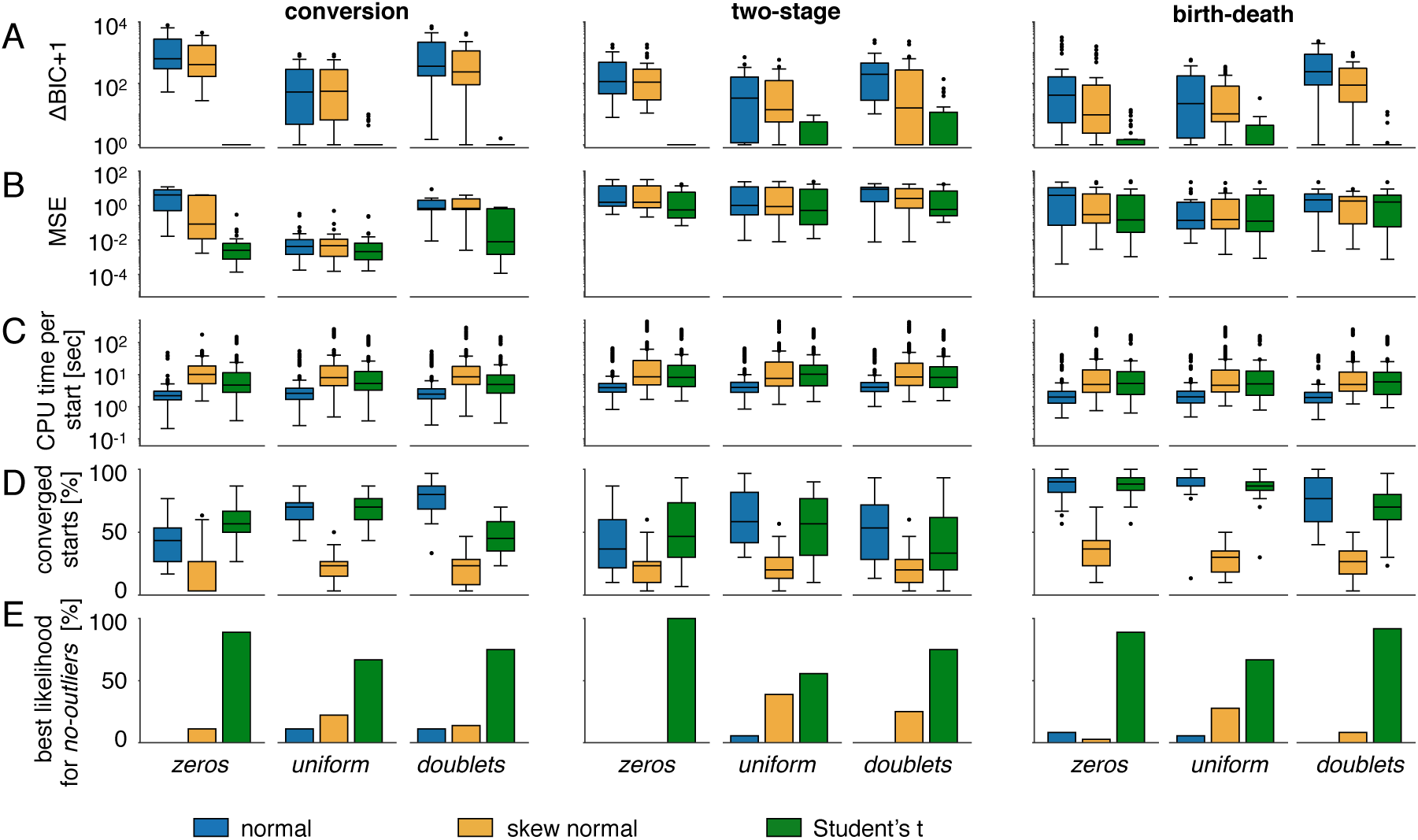
Results for the simulation study for the individual outlier scenarios. Comparison of (A) ΔBIC values, (B) MSE, (C) CPU time per optimization start and (D) number of converged starts for the distribution assumptions for the motifs conversion process, two-stage gene expression and birth-death process. Each boxplot in (C) has 1080 points (36 data sets and 30 optimization runs) and the other boxplots (A, B, D) comprise 36 points. (E) The likelihood values were calculated for the *no-outlier* data set using the parameters obtained for the outlier-corrupted data sets. The bars show how often the corresponding distribution provided the best likelihood value. A higher percentage indicates robustness against outliers.

### D.3 Birth-death process

The birth-death process is described by the following reactions:

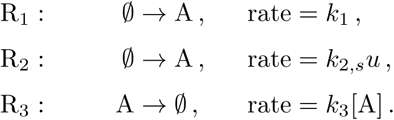

Reactions R_1_ and R_2_ describe the stimulus-independent and stimulus-dependent production of A and reaction R_3_ its degradation.

### D.4 NGF-induced Erk1/2 signaling

We used the model proposed in Loos et al. (2018), assuming cell-to-cell variability in total Erk1/2 levels and inter- and intra-subpopulation variability in cellular TrkA activity. The moments of the system were obtained using the sigma-point approximation. The whole data set with the model fits is shown in Fig. D2.

**Figure D2:**
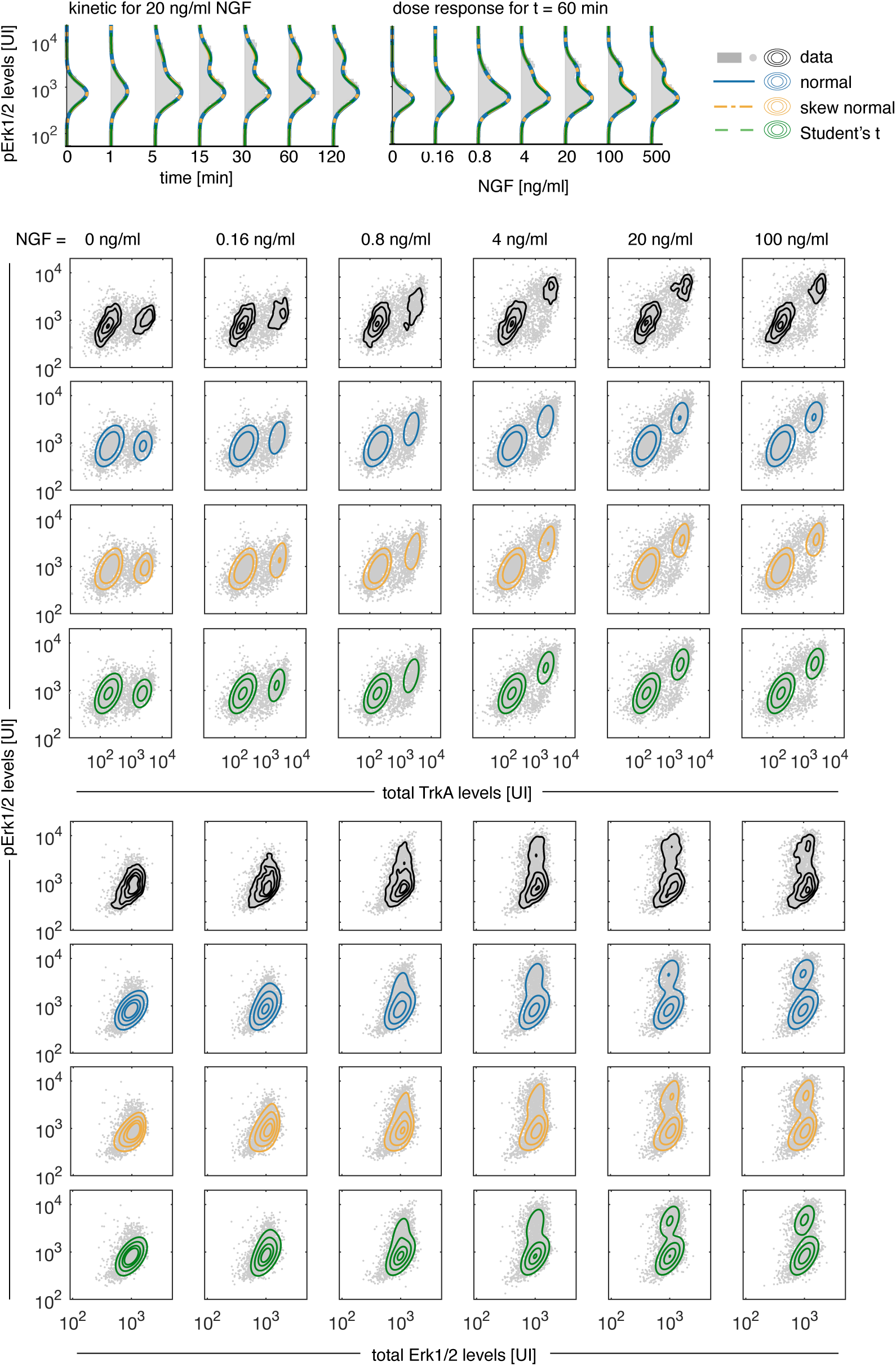
Data and model fits for NGF-induced Erk1/2 signaling. pErk1/2 kinetics and dose responses, and multivariate measurements of pErk/TrkA levels and pErk/Erk levels. The upper rows illustrate the data together with a kernel density estimate. The bottom rows visualize the data together with the contour lines of the hierarchical models.

